# *Bacillus subtilis* spore induces efficient generation of memory T cells via ICAM-1 expression on dendritic cells

**DOI:** 10.1101/2020.01.20.912584

**Authors:** Lulu Huang, Jian Lin, Yuchen Li, Penghao Zhang, Qinghua Yu, Qian Yang

## Abstract

The intestinal mucosa is the primary exposure and entry site of infectious organisms. Tissue-resident memory T cells (Trms) is an important first line of defense against infection in mucosal tissues, their function in intestinal immunization remains to be investigated. Here, we reported that the levels of local mucosal and systemic immune responses were enhanced through oral immunization with H9N2 whole inactivated virus (H9N2 WIV) plus spore. Subsequently, H9N2 WIV plus spore led to the generation of CD103^+^CD69^+^ Trms, which was independent of circulating T cells during the immune period. Meanwhile, we also found that *Bacillus subtilis* spore can stimulate Acrp30 expression in 3T3-L1 adipocytes. Moreover, adipocyte supernatant or spore also upregulated intercellular adhesion molecule-1 (ICAM-1) expression on dendritic cells (DCs) (*P*<0.01). Furthermore, the proportion of HA-tetramer^+^cells was severely curtailed when ICAM-1 expression was suppressed, which was also dependent on HA-loaded DCs. Taken together, our data demonstrated that spore promoted the immune response by stimulating Trms, which were associated with activation of ICAM-1 in DCs.

**Author summary:** Taken together, Bacillus subtilis spore combined with H9N2 WIV enhanced the mucosal antibody response and induced efficient intestinal-resident memory T cells. Then we demonstrated that spore can induce memory T cell formation through an ICAM-1-mediated contact of a dendritic cell (DC)-derived mechanism. Further, our findings indicated that Acrp30 from adipocytes induced by spore might increase ICAM-1 expression on DCs, which might provide new insight into the significance of adipocyte metabolism related molecules in regulating immunological memory T cells.

## Introduction

H9N2 subtype avian influenza virus, a low-pathogenicity avian influenza (AIV), has become endemic and pose a significant threaten to human and animal [1, 2], which can replicate in avian guts and spread by fecal-oral transmission [3, 4]. Mucosal immune is an effective way to block the infectious organisms through eliciting memory T cells against pathogens. Hence, acquisition of the protective immune responses at mucosal sites is a priority vaccination strategy to prevent pathogenesis. Although expert in eliciting immune responses, oral immunization with vaccination also induces poor mucosal effect when immunized with inactivated virus alone [5, 6]. Thus, urgent vaccine development involves inducing protective immune responses against potential pathogens on mucosal surfaces, suggesting a critical need for more efficacious adjuvants to improve vaccine potency. *Bacillus subtilis* spore, acts as an adjuvant, can strongly induce immune responses against pathogen, especially in modulating intestinal mucosal immunity through evoking tissue-resident memory T cell (Trm) [7–9].

Previous studies on *Bacillus subtilis* spore found that it can stimulate the secretion of cytokines as innate immune signaling, which is indispensable for efficient induction of adaptive immune responses during primary immunization [10]. Recent study found that mucosal immunization with Spore-FP1increased CD69^+^CD103^+^ Trm in the lung parenchyma [11]. Meanwhile, our previous study also confirmed *Bacillus subtilis* spore, as advantages of mucosal delivery, could regulate memory T cells in the intestine of piglets [12]. As we known, T cells is important for cell immunity, while vaccine-mediated intestinal T cell responses reveal a requirement for the addition of adjuvant for evoking a robust Trm response[13]. Recent reports have demonstrated that the expression patterns of lymph node homing receptors CCR7 and CD62L are closely related to the functional status of central memory T cells (Tcms) and effector memory T cells (Tems) [14]. Moreover, Trms mediate rapid clearance of and heterosubtypic protection against secondary IAV infections in mice [15, 16]. Further analyses have revealed that Trms were also detected in the intestinal mucosa, leading to tissue-specific influences [17–19]. Until recently, Trms, which express high levels of C-type lectin CD69 and low levels of the sphingosine-1-phosphate (S1P) receptor S1PR1, are thought to be phenotypically and functionally distinct from circulating memory T cells [20]. Furthermore, establishment of long-term and resident memory depends on the maintenance of CD103 and CD69 expression in T cells [21]. Thus, the features and processes of memory cells are involved in the retention and persistence of T cells in mucosal tissue, thereby promoting long-term protection for viral clearance [19].

Our study provided further insight into the potential immunopotentiator ability of *Bacillus subtilis* spore to assist PEDV WIV in the potentiation of immunity by upregulating memory T cells via oral immunization in piglets [12]. However, the specific mechanism in memory cell formation remains to be further studied. Previous studies suggest that intercellular adhesion molecule-1 (ICAM-1) is critical for establishing memory T cells following acute infection [22]. In addition, a substantial number of liver-resident memory populations are regulated by LFA-1-ICAM-1 interactions following LCMV immunization [23]. Lipid metabolism related molecules play important role in regulating ICAM-1expression [24]. Hence, our study try to illustrated the underline mechanism of whether *Bacillus subtilis* spore induce memory T cell formation through activating ICAM-1, as well as inducing Acrp30.

## Results

### Spore recruited and activated DCs

DCs are essential for the generation of T-cell immunity after mucosal immunization [25]. To investigate whether spore had the capacity to recruit submucosal DCs to form transepithelial dendrites (TEDs) for viral capture, we assessed TED formation in vitro and in vivo. Initially, in a DC/epithelial cell (EC) coculture system (Fig. 1A), we observed spore, but not medium, induced DCs to form TEDs across ECs at 30 min in cross-sectional images (Fig. 1B). Then, Ligated loop experiments at 0.5 h after spore adminstrated found DCs were apparently gathering to the laminal propria of ECs, which were significantly increased by spore compared with the control (Fig. 1C). Moreover, we found spore had the powerful capacity to increase the expression of CD40 and CD80 (CD80: *P* < 0.01, CD40: *P* < 0.01) (Fig. 1D) in coculture system, compared with medium alone. Furthermore, the release of proinflammatory cytokines (IL-1β, TNF-α) stimulated by spore, which also indicated the functional maturation of DCs (Fig. 1F) (*P* < 0.01). By the way, our result also observed spore significantly increased the length of DCs (Fig. 1E).

**Fig 1.**
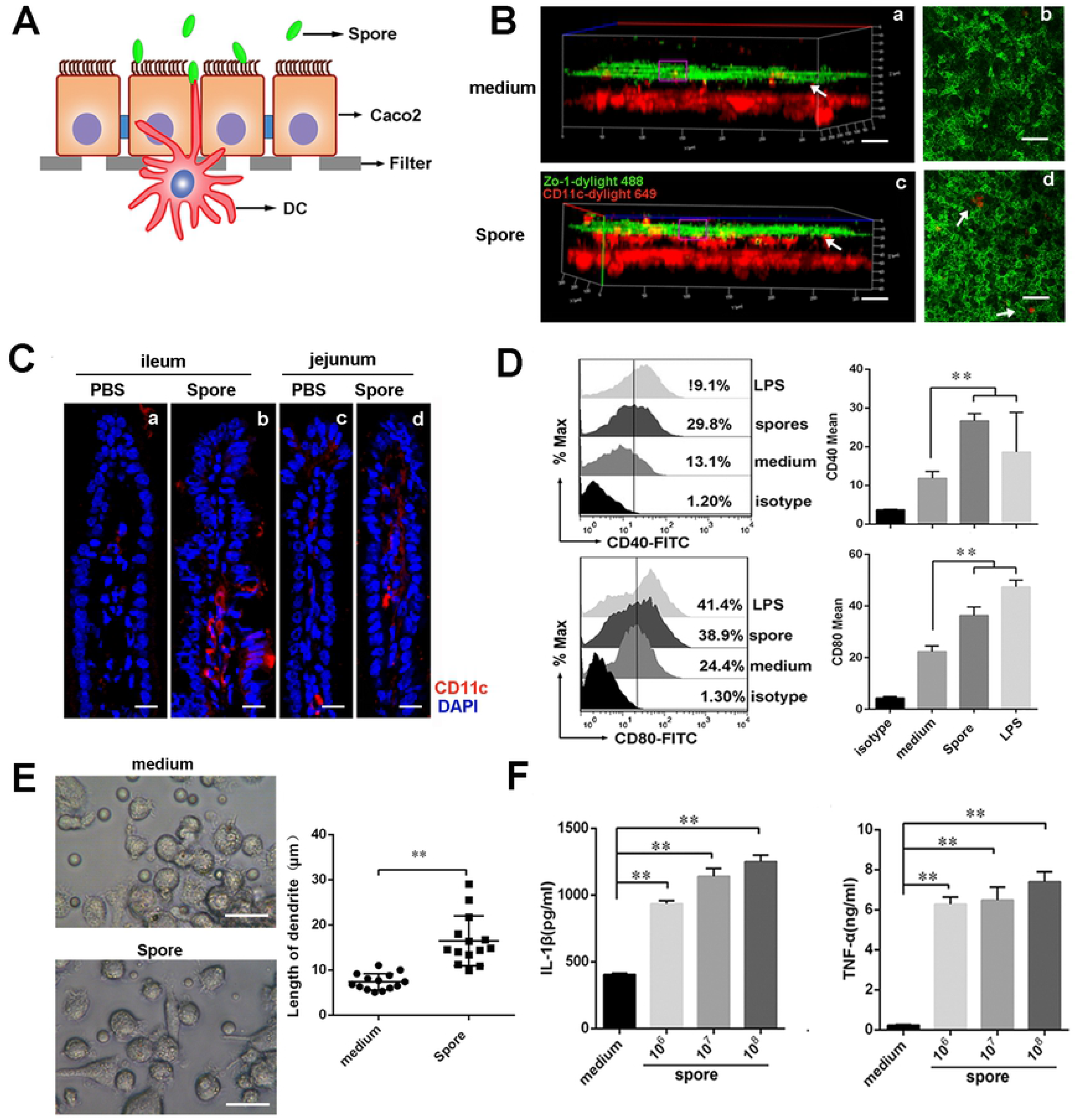
Spore activated dendritic cells *in vitro* and *in vivo*. (A) Schematic of the experiment used to study activation of DCs in the DC/EC coculture system. (B) In the coculture system, medium (a, b) and spore (c, d) were incubated on the apical side of the Caco-2 monolayer for 0.5 h. The filters were processed for immunofluorescence staining and observed using CLSM. A three-dimensional (3D) rendering of representative fields was obtained with ZEN2012 software. Submucosal DCs (CD11c, red) caused dendrites (white arrow) to creep through the tight junctions (TJs) of ECs (zo-1, green) in response to spore but not medium. Scale bars = 50 μm. (C) Ligated loops of mice were injected with spore *in vivo*, and intestines were isolated after 1 h and then processed for immunofluorescence staining. Cryosections stained with anti-CD11c antibody (red) and 4’, 6-diamidino-2-phenylindole (DAPI; blue) were observed under a confocal microscope. TEDs are indicated by arrows. Scale bars =10 μm. (D) In the coculture system, DCs were stimulated for 24 h with LPS (100 ng/ml) or spore (10^6^, 10^7^, and 10^8^ CFU/ml), and surface molecule expression on gated viable cells was measured by flow cytometry on gated viable cells. The phenotypic expression levels of CD40 and CD80 on DCs were analyzed by FACS. (E) DCs were treated with medium and spore separately for 24 h, and the morphology of DC dendrites was observed by microscopy. Scale bar = 20 μm. (F) Secretion of interleukin (IL)-1β and TNF-α in culture supernatants was measured by ELISA. The results are expressed as the mean ± SEM. Significance was tested against the unstimulated control by one-way ANOVA, \**P* < 0.05, \*\**P* < 0.01. One representative result of three similar independent experiments is shown.

### Spore facilitated H9N2 WIV to enhance H9N2-specific antibodies and induce T cell proliferation

Local secretion of IgA antibodies is the most important characteristic mediating oral adaptive immunity and mucosal protection. As shown in the immunization schedule, vaccine induced antibody responses were analyzed in the serum and mucosal fluids at different time points (Fig. 2A). Spore facilitated H9N2 WIV in enhancing the intestinal IgA response after oral immunization in mice (Fig. 2C). Similar changes in IgA levels in lung wash samples were also observed at 7 d, 35 d and 49 d (Fig. 2B) (*P* < 0.01), suggesting a marked effect of spore on mucosal responses in the lower respiratory tract. In addition, a trend reflecting increased levels of H9N2-specific IgG induced by spore plus H9N2 WIV was observed. Spore was able to significantly enhance the levels of IgG in the serum compared with PBS. As shown by the results, the levels of IgG at 21 d, 35 d and 49 d (Fig. 2D) (*P* < 0.01), IgG1 at 35 d (Fig. 2E) (*P* < 0.01) and IgG2a at 21 d, 35 d and 49 d (Fig. 2F) (*P* < 0.01) induced by H9N2 WIV plus spore were significantly higher than the levels induced by H9N2 WIV alone. In addition, serum collected from different groups of mice at 21 d and 49 d showed a powerful ability to inhibit hemagglutination against 4-HA units of H9N2 compared with antigen alone (Fig. 2I).We isolated lymphocytes from the spleen and mesenteric lymph node (MLN) 21 d post-immunization, and cells were restimulated with H9N2 WIV *in vitro*. We found that the proliferative index in MLNs was markedly increased in the spore plus H9N2 WIV group compared with that in the antigen-alone group (Fig. 2G) (*P* < 0.01). Similarly, the proliferative index in the spleen was increased (Fig. 2H) (*P* < 0.05), reflecting effective induction of systemic and local immune responses in mice.

**Fig 2.**
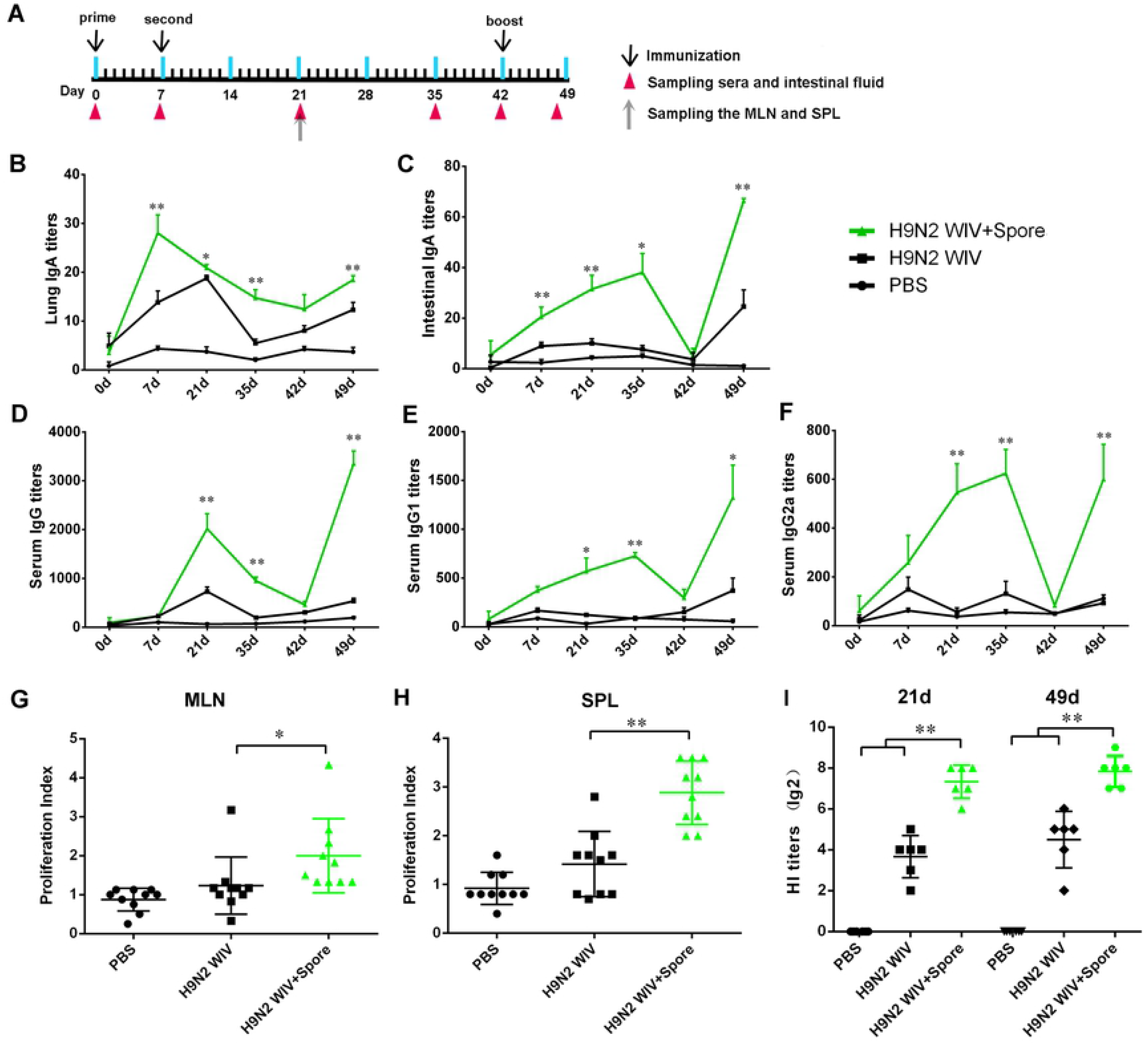
Spore facilitated H9N2 WIV to enhance antigen-specific antibodies. (A) Schematic of oral immunization and the sampling schedule of intestinal fluids, lung wash fluids, serum, mesenteric lymph nodes (MLNs) and the spleen. H9N2 WIV (20 μg) and spore (10^8^ CFU/ml) were orally administered to each mouse. Primary and secondary immunizations were performed at 0 d and 7 d, respectively. Booster immunizations were administered at 42 d. The details of the immunization schedule indicate the time point of immunization (black arrows above the line). Sampling is indicated by the time points of the serum, intestinal fluid and lung washing buffer collection arrows (below the line). (B-F) H9N2 specific IgA and IgG antibodies in mice post-immunization were determined by ELISA. Antigen-specific serum (D) IgG titers, (E) IgG1 titers, (F) IgG2a titers, and (B) mucosa IgA titers in intestinal wash and (C) lung wash fluids were detected at different time points. The asterisks indicate significant differences between H9N2 WIV plus spore and H9N2 WIV alone. (G and H)MLNs (G) and splenic (H) lymphocytes from immunized mice were isolated and restimulated with H9N2 WIV (10 μg/ml) *in vitro*. The proliferative response was detected by a CCK8 assay. (I) Hemagglutination inhibition (HI) titers were detected at 21 d and 49d. The results are expressed as the mean ±SEM. *P* values < 0.05 were considered to be statistically significant (\**P* < 0.05, \*\**P* < 0.01) (n=6).

### Spore-adjuvanted immunization induced CD69^+^CD103^+^ Trms in intestinal tissue

To further evaluate whether spore could induce Trm formation *in vivo*, we performed oral immunization in mice with PBS, H9N2 WIV alone or H9N2 WIV combined with spore. In the present study, spore-adjuvant immunization significantly upregulated the expression levels of the Tcm surface makers CD62L and CCR7 in blood at 7 d after primary immunization (S1 Fig. A) (*P* < 0.01). Nevertheless, no significant difference was observed at 45 d (S1 Fig. C). Recently, Trms were found in several tissues, including the intestinal mucosa [26] and lung [27]. Since spore could not cause the development of Tcms in blood following immunization, we next examined presence and proportion of Trms in the intestine. Two surface markers, CD103 and CD69, have been considered in distinguishing Trms from other memory T cells [28–30]. Intestinal tissues were harvested from immunized animals, and CD3-positive cells were then assessed for the expression of the tissue retention markers CD69 and CD103 at 7 d, 14 d and 45 d (Fig. 3A). No effect on the frequency of CD69^+^ CD103^+^ cells among CD3^+^ T cells was observed at 7 d (Fig. 3B). Notably, flow cytometry analysis showed that spore plus H9N2 WIV induced 21.5% CD69^+^CD103^+^ Trms at 14 d (Fig. 3C) and 49.1% CD69^+^CD103^+^ Trms at 45 d (Fig. 3D). Furthermore, the expression of IFN-γ^+^ T cells markedly increased at 45 d (Fig. 3E) after H9N2 WIV restimulated IMALs. These data supported the capacity of a mucosal vaccine to induce substantial T cell responses after oral immunization.

**Fig 3.**
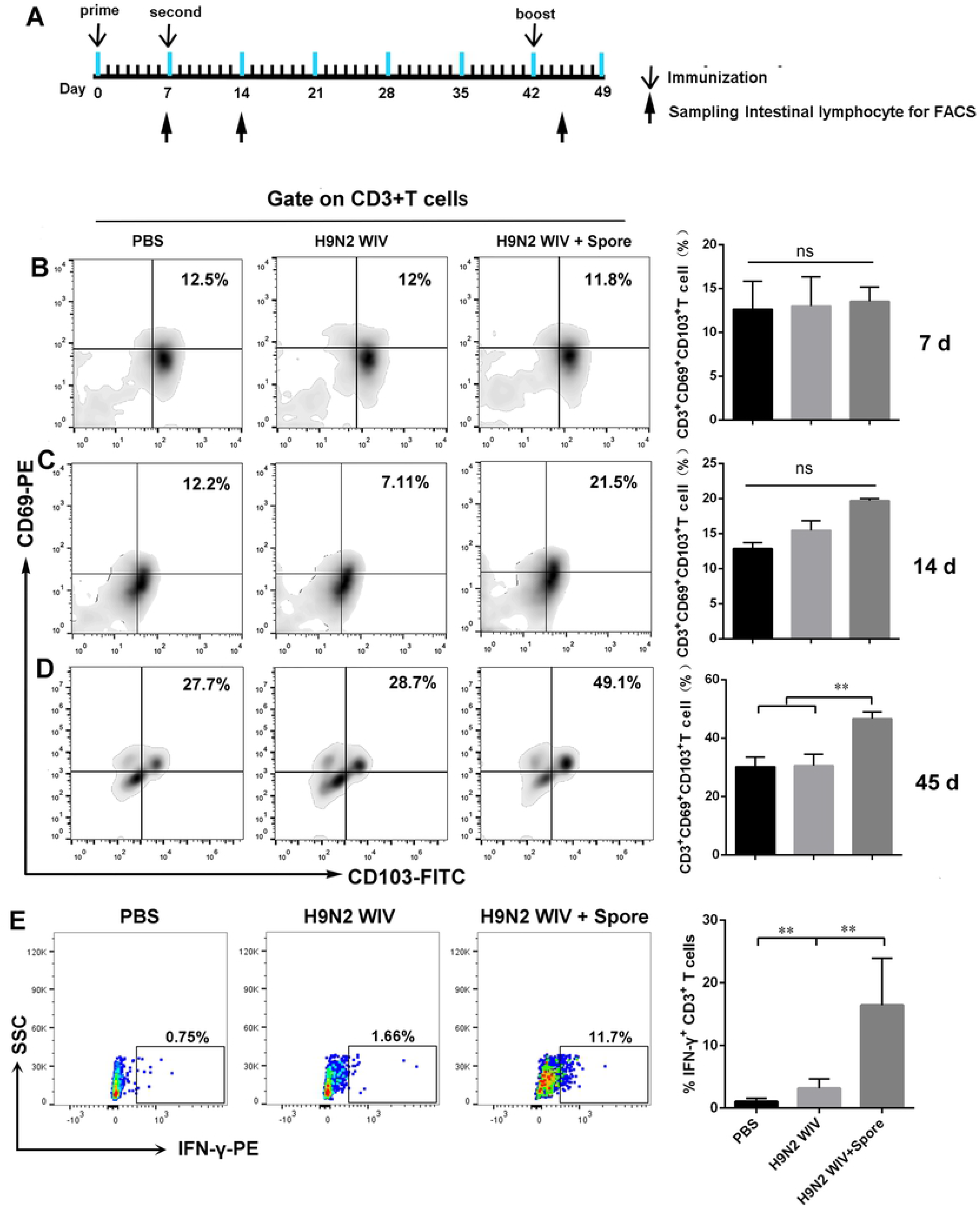
The frequency of CD69^+^CD103^+^ Trms after oral immunization in the intestinal tract. (A) Schematic experimental design to examine the frequency of Trms after oral immunization with different vaccines. (B-D) The frequency of Trms (CD3^+^ CD69^+^ CD103^+^) was detected in the intestinal mucosa at 7 d (B), 14 d (C) and 45 d (D) after priming immunization. A gating strategy was applied in this study to determine the memory cell phenotype of CD3^+^ T cells according to CD69 and CD103 from in intestinal tissue. (E) IFN-γ expression in CD3^+^ T cells from immunized mice was detected by FACS following H9N2 WIV recall. Data are represented as the mean ± SEM (n=6). \**P* < 0.05, \*\**P* < 0.01. One representative of two similar independent experiments was shown.

### Spore-adjuvanted immunization induced HA-specific Trms in intestinal tissue

We also investigated antigen-specific Trms after mucosal immunization. To detect whether FTY720 treatment could inhibit lymphocyte circulation, FTY720 was administered to inhibit the circulation of T cells six weeks after the primary vaccination, as illustrated in Figure 4A. In addition, mice were injected i.v. with anti-CD45-FITC antibodies 10 min prior to harvesting blood and intestinal Peyer’s patches (PPs). The vast majority (> 99.9%) of the intestine failed to be stained by anti-CD45 antibodies after treatment with FTY720 (S2 Fig). Interestingly, the influenza HA-specific T cells following mucosal immunization were significantly increased in H9N2 WIV plus spore-treated mice and were not altered by FTY720 treatment (Fig. 4C and D) (*P* < 0.05). Thus, we can speculate that these memory cells also show a bias toward tissue residency. Consistent with the flow cytometry results, we also observed accumulation of influenza HA-specific cells in the intestinal tract, and imaging using microscopy showed that at 5 weeks after oral immunization, allophycocyanin (APC)-tetramer^+^ T cells were readily detectable in the PPs of the ileum (Fig. 4B). Together, these results demonstrated that mucosal immunization with H9N2 WIV plus spore can generate influenza-specific T cells in the intestinal tract.

**Fig 4.**
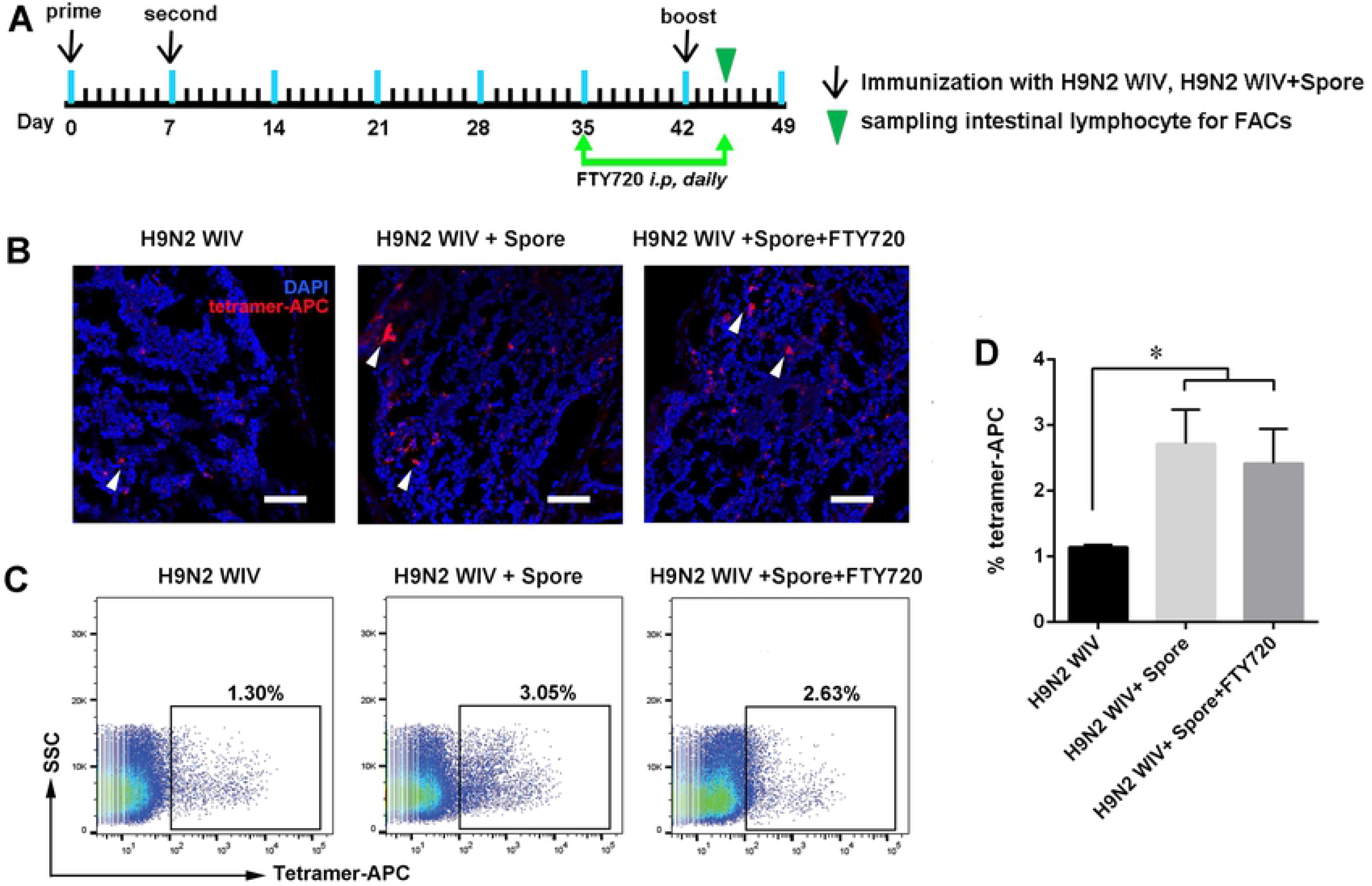
Evidence of HA-specific T cells after oral immunization in intestinal tract. (A) BALB/c mice were immunized as described previously, and lymphocytes from the intestine were analyzed by HA-specific tetramer staining at 45 d to detect the effect of FTY720 treatment on T cells. Five weeks after immunization, immunized mice were treated with FTY720, 1 mg/kg by i.p. daily for 10 d. (B-D) APC-tetramer^+^ T cell populations were analyzed and compared with those of immunized mice treated with H9N2 WIV plus spore but not treated with FTY720 by FACS (C, D) and confocal microscopy (B). The scale bar represents 50 μm. Data are represented as the mean ± SEM (n=6). \**P* < 0.05, \*\**P* < 0.01. One representative result of two similar independent experiments is shown.

### Spore upregulated ICAM-1 expression after mucosal immunization

Upon revealing the activation of DCs, we then assumed molecules that could be regulated in the intestinal microenvironment. To this end, a proteome profiler array analysis was performed to measure 110 proteins from the lysates of the intestinal tract at 7 d or 45 d after oral immunization of mice with PBS or spore (Fig. 5B). In particular, ICAM-1 and adiponectin/Acrp30 was relatively upregulated in intestinal tissue after spore immunization compared with PBS treatment (Fig. 5C). Based on these considerations, we focused on ICAM-1 and its potential action on DCs. High expression of ICAM-1 was also verified in intestinal tissue by western blot and immunohistochemical staining (Fig. 5D and 5E) (*P* < 0.01).

**Fig 5.**
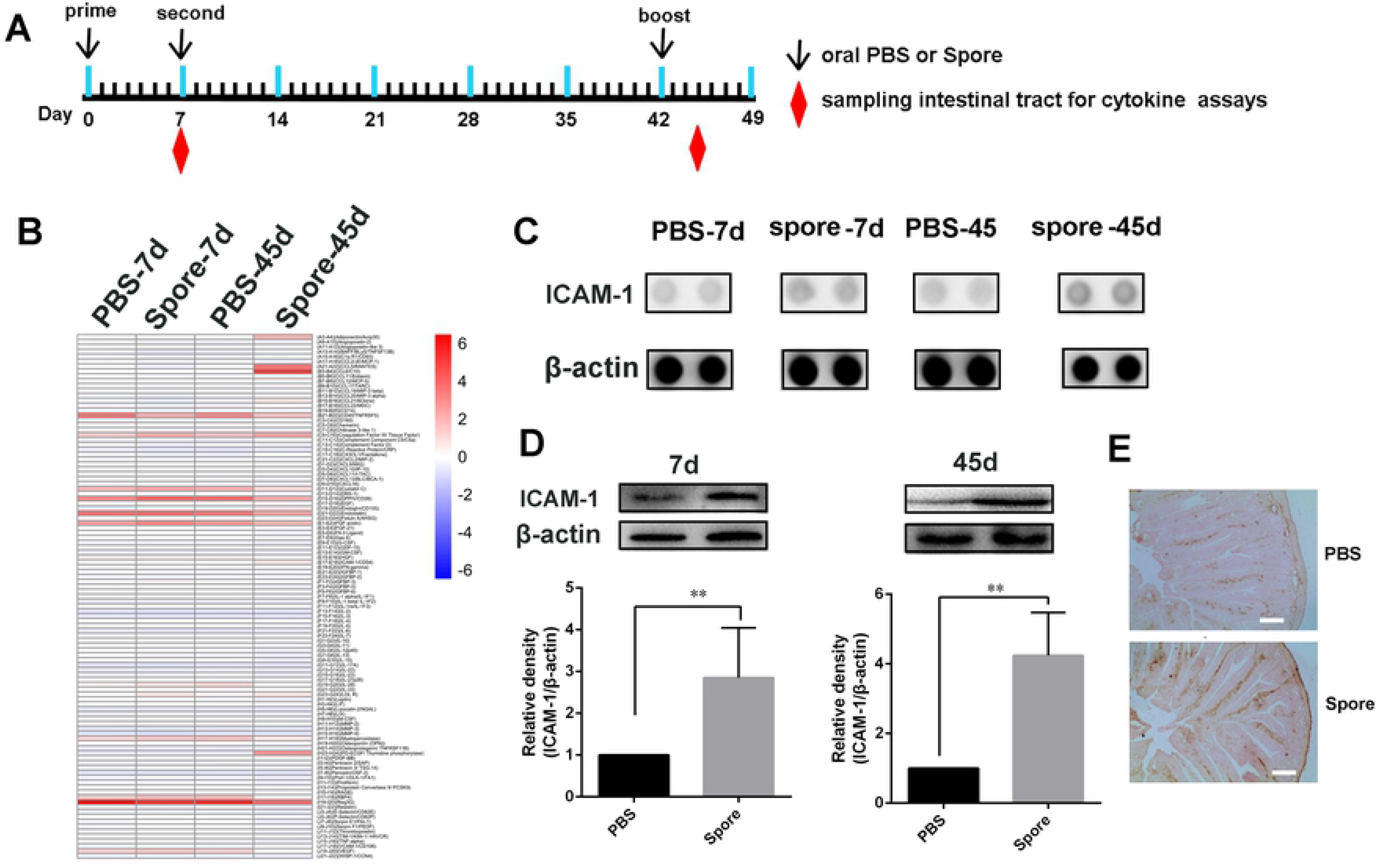
ICAM-1 expression in the intestinal tract after oral administration of spore. (A) Schematic experimental design of oral immunization with PBS or spore and the sampling schedule of intestinal tissue and lymphocytes. (B) Gray value intensity was detected utilizing chemiluminescence and membranes can be assessed for protein levels. Intensity is shown in a pseudocolor scale (from low [blue] to high [red]). (C) Mouse intestinal whole-tissue lysate was analyzed by an XL mouse antibody array. The solid black circles indicate proteins secreted by mice. (D) Western blot analysis revealed the time-dependent upregulation of ICAM-1 in the ileum tissue following immunization with spore at 7 d and 45 d. Equal proteins loading was confirmed using the house-keeping gene β-actin. (E) Induction of ICAM-1 expression was confirmed by immunohistochemistry (IHC) staining in the ileum. The scale bar represents 20 μm. Data are represented as the mean ± SEM (n=6). \**P* < 0.05, \*\**P* < 0.01. One representative of three similar independent experiments is shown.

### ICAM-1 expression on DCs increased after oral spore treatment

Previously, it has been showed that upregulation of ICAM-1 on APC could regulate the generation of central memory cells [31]. According to previous results, we showed that ICAM-1 expression was significantly increased in DCs following spore treatment (10^6^, 10^7^, 10^8^ CFU/ml) depending on the concentration (Fig. 6A and B) (*P* < 0.01). Moreover, this finding suggested that the effect coincided with the increased levels of ICAM-1 measured by qPCR and IF (Fig. 6C and D) (*P* < 0.01). Our current results implied that spore first stimulated DC recruitment to the LP *in vivo* (Fig. 1) and then activated ICAM-1 molecule to further stimulate T cells. To further define whether this phenomenon was reproducible *in vivo*, we performed FACS for ICAM-1 expression in CD11c^+^ DCs. We noted that ICAM-1 was upregulated in the intestinal submucosal DCs of mice after immunization with spore plus H9N2 WIV (Fig. 6E). Furthermore, the number of ICAM-1^+^ DCs (CD11c^+^) was detected in the intestine using the double fluorescence staining method (Fig. 6F). In brief, ICAM-1 expression was notably increased on DCs after spore stimulation *in vitro* and *in vivo*.

**Fig 6.**
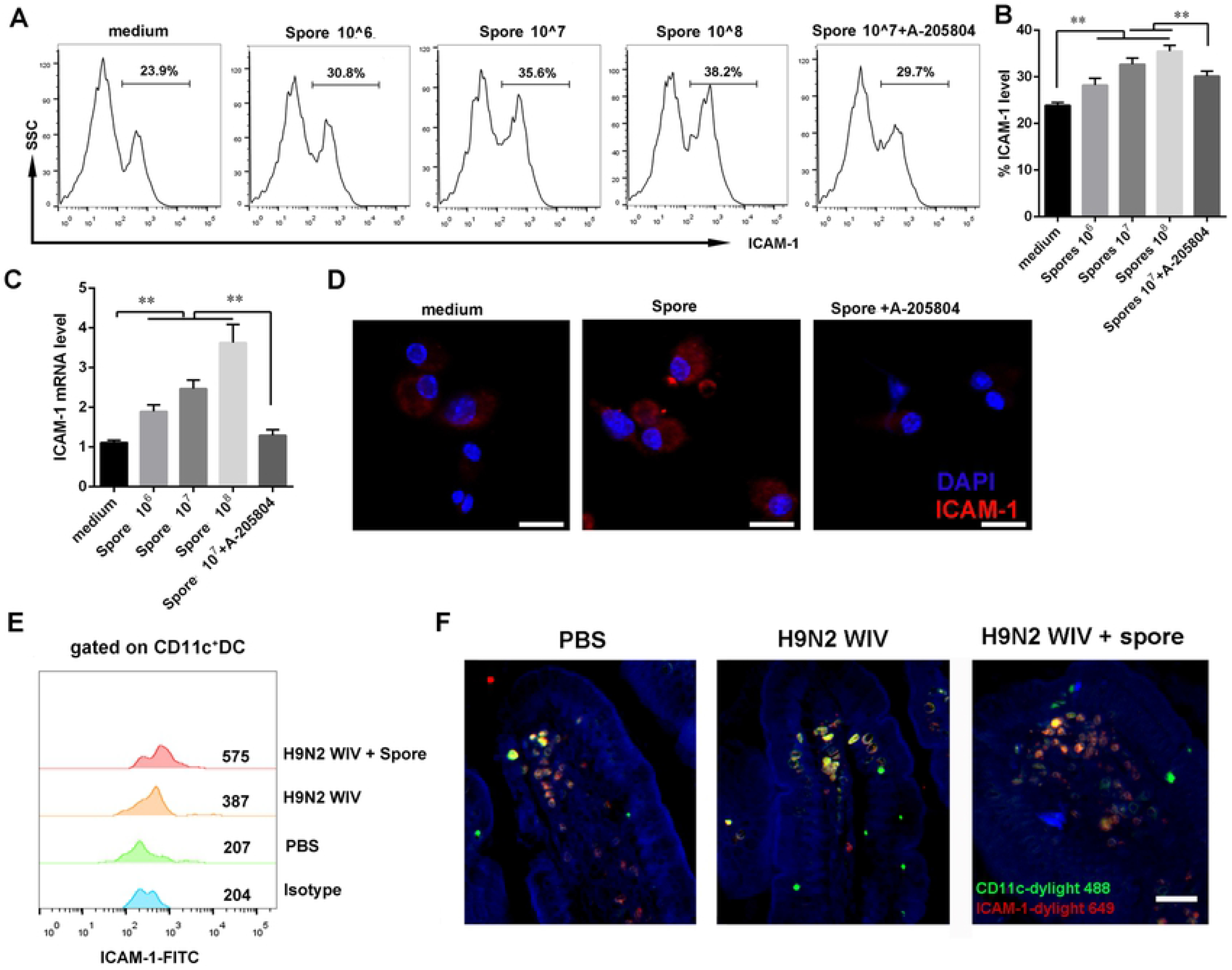
ICAM-1 expression on DCs was increased after stimulation with spore. DCs were treated with spore (10^6^ CFU/ml, 10^7^ CFU/ml and 10^8^ CFU/ml) or spore plus A-205804 (10 μM) for 24 h. (A, B) ICAM-1 expression was detected on DCs after spore treatment by FACS. (C) ICAM-1 mRNA expression was measured by RT-qPCR and confocal microscopy. (D) ICAM-1 protein expression was detected by immunofluorescence after DCs were incubated with medium, spore or spore plus ICAM-1 inhibitor A-205804 for 24 h. The scale bar represents 20 μm. (E) Cells were collected from the intestinal tract, and the MFI of ICAM-1 gated from CD11c^+^ DCs was detected by FACS. (G) ICAM-1^+^ CD11c^+^ double positive cells in the intestine were strong positivity stained by immunofluorescent staining. The scale bar represents 20 μm. Data are represented as the mean ± SEM (n=6). \**P* < 0.05, \*\**P* < 0.01. One representative result of three similar independent experiments is show

### Spore stimulated Acrp30 expression in 3T3-L1 Adipocytes

Since we have demonstrated that spore could enhance the Acrp30 level in intestine (Fig. 5A). In order to determine stimulation effects of spore treatment on differentiated 3T3-L1 cells, we performed the induction culture assay of adipocytes. Initially, cells displayed a fibroblast phenotype. Then, cell morphology changed and cells accumulated lipid droplets internally during the process of differentiation. After 12 days, when the adipocytes were mature, almost the entire cell volume was stained red by red oil (Fig. 7A). We subsequently determined the effects of spore on Acrp30 protein expression. Spore at 10^6^ CFU/ml was sufficient to elicit up-regulation of the Acrp30. Treatment of 10^7^ CFU/ml of spore led to a 2-fold increase in the Acrp30 protein amount in comparison with the control without spore (Fig. 7B). The extent of spore-induced increase in protein levels was positively associated with the concentrations of spore, indicating that there is a dose-dependent effect of spore on the increase of adiponectin proteins. We also investigated whether Acrp30 induced by spore could increase ICAM-1 expression on DCs. The cultural supernatant from differentiated 3T3-L1 cells treated by spore or PBS could stimulate the ICAM-1 expression on DCs (Fig. 1C and D). Our data demonstrated that spore effectively up-regulated the expression of the adiponectin and enhanced ICAM-1 expression on DCs.

**Fig 7.**
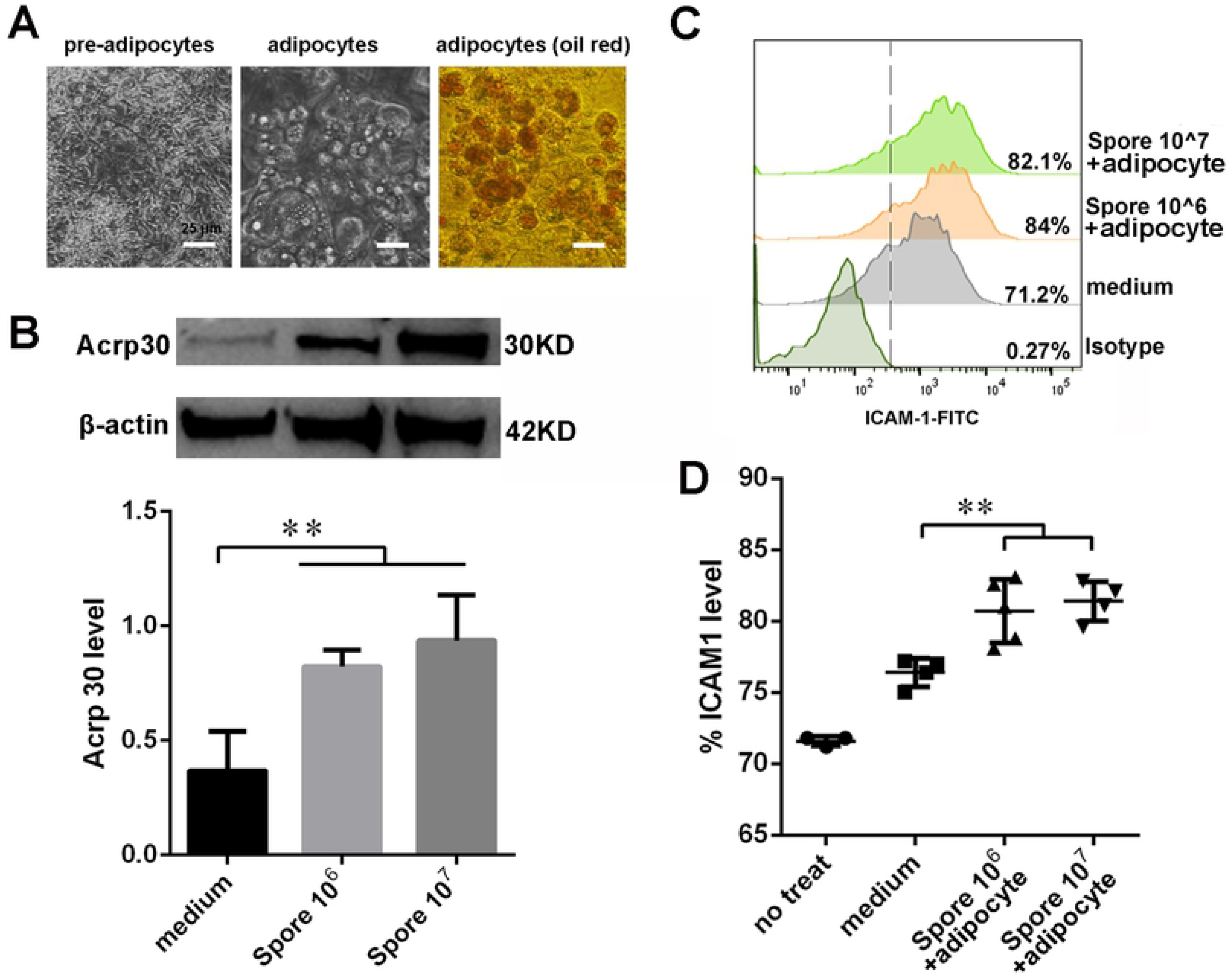
Secretion of Acrp30 from adipocyte induced by Spore increased ICAM-1 expression on DCs. (A) 3T3-L1 cells were differentiated into adipocytes according to the differentiation protocol. Phase contrast images of 3T3-L1 cells from 0 day induction (pre-adipocyte) to ten days post-induction (adipocyte). Triglyceride staining of 3T3-L1 cells with Oil Red O. (B) Differentiated 3T3-L1 cells were treated with spore for 24 h. Western Blot assay was performed to detect the levels of adiponectin in the cells. The bar graph represents quantification of the relative protein levels of adiponectin. (C-D) DCs were treated with medium from Differentiated 3T3-L1 treated with spore for 24 h. Flow cytometry analysis was performed with anti-ICAM-1-FITC staining. A representative blot is shown in the upper panel. Data are represented as the mean ± SEM. \**P* < 0.05, \*\**P* < 0.01. One representative of three similar independent experiments is shown. Bars: 25 μm.

### ICAM-1-dependent DCs induced the generation of HA-specific T cells

Pretreatment with spore and/or S-205804 suppressed the ICAM-1 expression of DCs or anti-ICAM-1 neutralizing antibody, and the CD44^+^ CD69^+^ phenotype markers of T cells were detected by FACS. As expected, flow cytometry analysis showed significantly increased levels of the CD44^+^CD69^+^ phenotype markers of Trms in DCs with spore treatment, and the treatment of ICAM-1 inhibitor A-205804 suppressed the CD44^+^CD69^+^ phenotype induced by spore (Fig. 8A and B)(*P* < 0.01).

**Fig 8.**
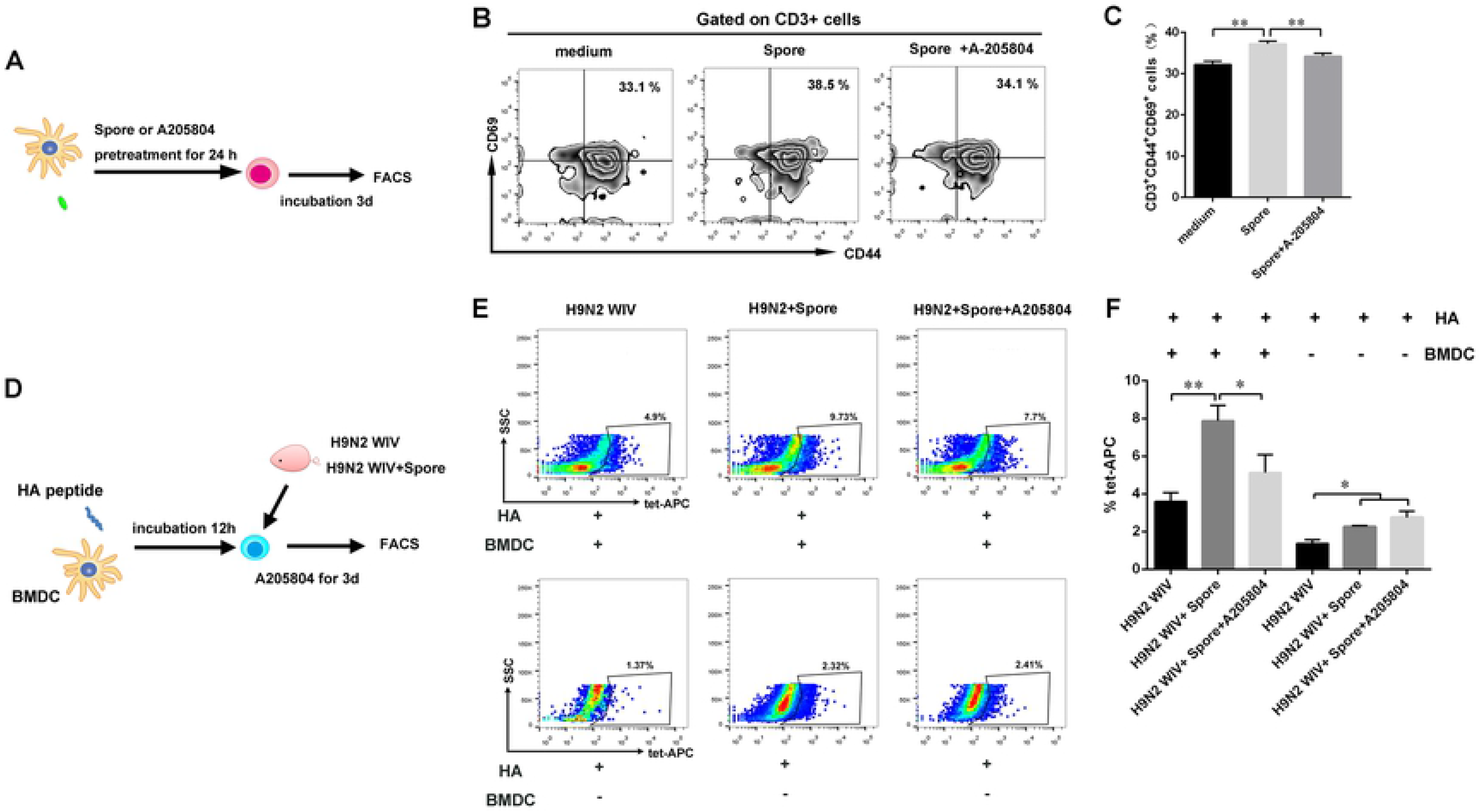
ICAM-1 dependent in DCs increased HA-specific Trm formation. (A) DCs were pretreated with spore and/or A204804 and then co-incubated with sorted CD3^+^ T cells from wild-type mice for 3 d. (B, C) Gated T cells were analyzed for the surface markers of memory cells CD44 and CD69 by FACS. (D) Six weeks after the primary vaccination, the intestines were dissected to prepare lymphocytes. BMDCs (5 × 10^5^ cells/well) were stimulated with HA_518-526_ (IYSTVASSL) peptide (10 μl) overnight. Antigen-pulsed DCs were used as APCs to stimulate IMALs (1 × 10^6^ cells/well) for 5 d. (E, F) Frequency of APC-tetramer^+^ T cells was detected by flow cytometry. Representative flow cytometry results and graphs for statistical analysis are shown. A representative result of two similar independent experiments is shown. Data are presented as the mean ± SEM. \**P* < 0.05, \*\**P* < 0.01.

**Fig 9.**
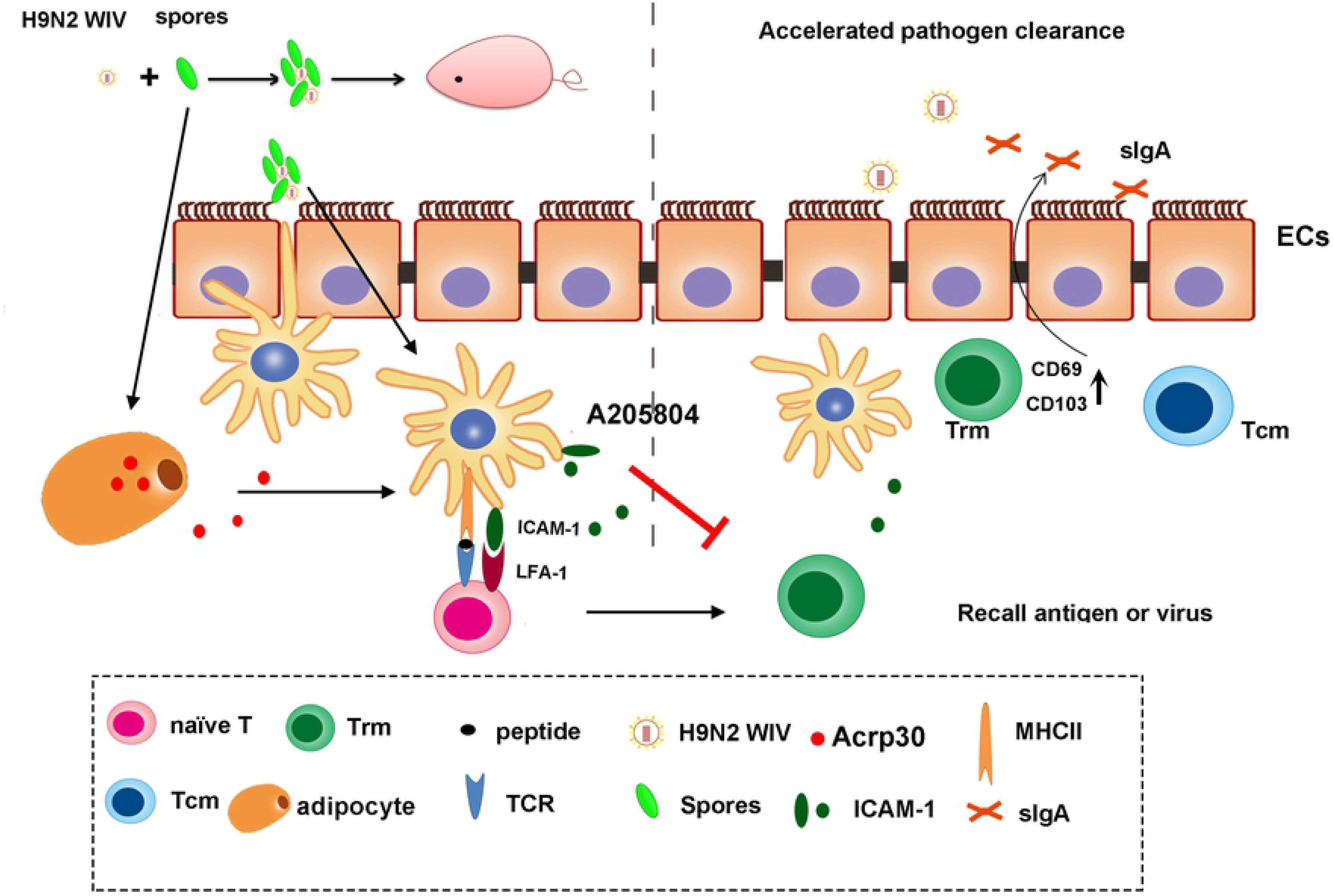
Overview of generating memory T cell generations after mucosal vaccination. Upon mucosal vaccination, DCs were stimulated with H9N2 WIV plus spore and then migrated to the draining lymph nodes and stimulated naïve T cells. Tcms recirculated between the blood and lymphoid organs or entered peripheral tissues. Vaccination also activated DCs to express ICAM-1 cytokines as well as LFA-1 binding to ICAM-1 on DCs. In addition, Acrp30 induced by 3T3-L1 differentiated adipocyte with the treatment of spore might generate the ICAM-1 expression on DCs. Then, ICAM-1 expression on DCs could upregulate proportion of antigen-specific Trms, which was altered after treatment of ICAM-1 inhibitor. The surface markers of Trms, such as CD69 and CD103, were upregulated after mucosal immunity. Local reactivation of mucosal Trm formation was induced by the ICAM-1 molecule, which triggered DCs recruitment and cytokine expression. The existence and maintenance of Trm subsets in intestine could accelerate pathogen clearance.

To assess whether HA-tetramer^+^ specific T cells persisted in the intestine at 6 weeks after immunization, antigen-loaded DCs with or without ICAM-1 inhibitor were incubated with IMALs isolated from immunized mice at 37°C for 3 d (Fig. 8D). Flow cytometric results showed that spore plus H9N2 WIV induced more HA-tetramer^+^ specific cells than H9N2 WIV alone at the present of antigen-pulsed DCs (Fig. 3E and F). Taken together, these results indicated that DCs partially require the expression of ICAM-1 for the generation of antigen-specific Trms, which were altered in the presence of ICAM-1 inhibitor added in DCs.

## Discussion

Oral immunization is beneficial for eliciting mucosal immune responses against pathogens that invade through a mucosal surface [32]. Here, we used a mucosal immunization strategy known as “prime-second-boost” and dissected the multifaceted adaptive immune mechanisms after mucosal immunization with spore plus H9N2 WIV. This strategy may represent an ideal platform for immunological protection and lead to robust antibody responses to generate long-lasting immunological memory.

This platform has numerous advantages as spore survived well and favorably stimulated DCs. Additionally, APCs such as DCs result in the generation of different fates for T cells as distinct populations of memory T cells in the absence of antigen [33]. Recent results prove that potent activation signatures in macrophages and bone marrow DCs with *Bacillus subtilis* spore treatment are accompanied by increased expression levels of the maturation markers CD40 and MHC classes I and II on DCs [34]. Moreover, the recruitment of DCs, particularly CD103^+^ DCs, promotes CD103 expression on immune cells and is essential for the efficient induction of Trms [35]. Subsequently, we focused on the innate immune system. Here, we performed experiments using a DC/EC coculture system and showed that spore played important roles in facilitating the delivery of DCs across intestinal mucosal barriers at an early stage of mucosal immunity and induced DC maturation, including the secretion of IL-1β. In accordance, mucosal expression of IL-1β is a sufficient and crucial mediator of Trm formation [36].

ICAM-1 expression is known to play a crucial role in the proper generation of T cell memory responses by APCs [37]. Furthermore, we observed that ICAM-1 had a much stronger impact on the initiation of mucosal memory T cells. Thus, the difference in mucosal T cell numbers and function between the immunized animals in the different groups likely reflects the response to spore-induced recruitment signals in the intestine. In this study, we identified ICAM-1 as a critical regulator of DCs in inducing memory T cell information. When the expression of ICAM-1was inhibited, the memory T cell phenotype markers CD44 and CD69 were observed less frequently than in the regular DC group. Current studies have shown that potential KLRG1-CD8^+^ Trms precursors with increased expression of CD69 can be isolated from the intestine, which remain the most reliable markers for intestinal-resident T cells in mice [16, 38]. A previous model predicted that ICAM-1 is required to augment the priming process, likely by promoting the recruitment of naïve T cells, prolonging cell-cell interactions, facilitating cytokine signaling, and permitting the differentiation of memory T cell precursors [22]. However, mucosal cytokines such as IL-1β induced by spore were not sufficient to induce differentiation of antigen-experienced T cells into Trms, which indicated the requirement for ICAM-1 upregulation on DCs in mucosal tissues. Our findings indicated that Acrp30 from adipocytes induced by spore increased ICAM-1 expression on DCs, which might provide new insight into the significance of adipocyte metabolism related molecules in regulating immunological memory T cells.

To investigate immunological antibodies associated with mucosal responses, we first evaluated antigen-specific antibody production in serum and mucosal fluid. We found that mice immunized with spore plus H9N2 WIV produced more specific IgG in the serum and IgA in the mucosal fluid compared with mice treated with H9N2 WIV alone. A similar trend was observed for lymphocyte proliferation after recalling antigens. Furthermore, we demonstrated that SF produced by *Bacillus subtilis* spore also induced more antibodies through the Th2 response, thus differing from spore. Overall, these findings may reflect a detection limitation in our assays or at least some potential metabolites of *Bacillus subtilis* spore as active components of mucosal immune enhancement.

Here, spore was able to induce a dramatically larger percentage of proliferating lymphocytes, indicating either a higher frequency of memory cells or cells with a higher proliferative capacity at the very least. Along with conventional T-cell activation signatures, we also observed a striking accumulation of gross CD69^+^CD103^+^ Trms in intestinal tissue after immunization with spore plus H9N2 WIV. Factors necessary for the establishment of Trms may play critical roles in this process and are not well understood. Recent vaccine studies have demonstrated that mucosal administration of antigen is important for the establishment of localized T cell responses [16]. In an optimally immunized individual, Trms were more protective than circulating memory cells based on their location and function [39]. Regarding the reason why no Trms are directed against spores themselves, spores may act as a mammalian commensal agent [40], thus suppressing the mobilization of Tefs that would lead to their clearance.

## Methods

### Animals and ethics statement

This study was approved by the Ethics Committee of Animal Experiments center of Nanjing Agricultural University. All animal studies were approved by the Institutional Animal Care and Use Committee of Nanjing Agricultural University (SYXK-2017-0007), and followed the National Institutes of Health guidelines for the performance of animal experiments. Specific pathogen-free C57BL/6 (4 to 6 weeks) and Balb/c (6 to 8 weeks) mice were obtained from Comparative Medical Center of Yangzhou University (Jiangsu, China). All animals were conducted at an animal facility under pathogen free conditions.

### Vaccine preparation

The influenza A/Duck/Nan Jing/01/1999 H9N2 virus was generously provided by the Jiangsu Academy of Agricultural Sciences [41]. The H9N2 virus was purified using a discontinuous sucrose density gradient. H9N2 WIV is normally inactivated via incubation at 56°C for 0.5 h to achieve a complete loss of infectivity. The *Bacillus subtilis* SQR9 strain was kindly supplied by Professor Shen of Nanjing Agricultural University [42].

### Immunogenicity study

Six-week-old Balb/c mice were orally immunized with H9N2 WIV (20 μg) alone or in combination with spore (10^7^ CFU) three times (at 0, 7 and 42 d). The mice were euthanized, and samples were collected at one-week intervals after the primary immunization. The levels of specific IgA in intestinal lavage fluid and specific IgG, IgG1, and IgG2a in serum were detected by ELISA. In brief, H9N2 WIV (2 μg/ml) antigens were coated onto a plate overnight, followed by blocking for 2 h with PBST containing 3% BSA. Intestinal lavage fluid and serum were diluted in PBS with 0.1% BSA and added to the plate in triplicate for 1.5 h at 37°C. Following five washes, HRP-conjugated rabbit anti-mouse IgG was incubated on the plate for 1 h. OD_450_ values were read on a Tecan plate reader at 450 nm absorbance. A hemagglutination inhibition (HI) test was performed according to a previously described procedure [43].

### Intestinal mucosa associated lymphocyte (IMAL) isolation

IMALs were isolated as described previously [44]. In brief, the intestine was opened longitudinally after the removal of residual mesenteric fat tissue. The tissue was then dissected into pieces and thoroughly washed with ice-cold PBS followed by digestion in 0.5 mg/ml collagenase D (Sigma), 0.5 mg/ml DNase I (Roche) and 50 U/ml dispase (Sigma) in Dulbecco’s phosphate buffered saline (DPBS) containing 5 mM EDTA, 4% fetal calf serum, and 100 μg/ml penicillin/streptomycin for 30 min at 37°C with slow rotation (100 rpm). After incubation, cells were collected and passed through a 70-μm strainer (BD Biosciences) and washed once with cold RPMI-1640. Then, the cells were resuspended in 6 ml of the 30% fraction of a Percoll gradient and overlaid on 6 ml of the 70% fraction in a 15-ml Falcon tube. Percoll gradient separation was performed by centrifugation at 300 g for 20 min. IMALs were collected at the interphase of the Percoll gradient, washed once, and resuspended in cold RPMI-1640 with 5% FBS. The cells were used immediately for experiments.

### General flow cytometry and cell sorting

For most experiments, cells were first stained with an Fc receptor blocker (1:20 dilution; eBioscience). For surface staining, cells were then stained with a mix of fluorescent antibodies in flow cytometry buffer for 30-45 min at 4°C for 0.5 h per the manufacturer’s guidelines. For Trm FACS, cells were stained with CD3-APC (145-2C11, eBioscience), CD103-FITC (2E7, eBioscience) and CD69-PE (H1.2F3, eBioscience) separately. For Tcms in blood, cells in 50 μl blood were stained with CD3-percp-cy5.5 (1452C11, Miltenyi Biotec), CD62L-APC (REA828, Miltenyi Biotec) and CCR7-PE (REA685, Miltenyi Biotec). Then, the whole blood was filtered through a 70-μm cell strainer, and the suspensions were incubated with an ammonium chloride potassium lysis buffer for 30 min at RT. For intracellular staining, the cells were incubated with 50 ng/ml phorbol myristate acetate (PMA; Sigma), 750 ng/ml ionomycin (Sigma), and 10 μg/ml brefeldin A (Invitrogen) in a cell culture incubator at 37°C for 5 h. After surface staining, the cells were resuspended in fixation and permeabilization solution (BD Biosciences) for 45 min at 4°C. Consistent with previous reports, the signature cytokines, interleukin (IL)-4 and IFN-γ, were measured on a BD FACS Verse and analyzed with FlowJo v.10.

For the lymphocyte enrichment assay, naïve CD4 or CD3 T cells were purified by negative selection similar to previously described methods [45]. Briefly, single-cell suspensions of lymph node (LN) or enteric cells were incubated with the following dilutions provided for each. For staining, a 100 μl volume of the antibody cocktail (BD Biosciences) was used per tissue sample from one mouse. Resuspended cells were incubated in an antibody cocktail for 15-30 min at 4°C in the dark. After washing the cells with 10 ml of PBS, the mixture was passed over a magnet following the manufacturer’s instructions. The purity of the flow-through fraction was routinely > 90%.

### Adipocyte differentiation culture

Mouse 3T3-L1 pre-adipocyte cells were cultured and differentiated as previously described [46]. Briefly, 3T3-L1 were grown in regular medium (high-glucose Dulbecco’s minimum essential medium (DMEM) supplemented with 10% FBS containing 1% penicillin and streptomycin). About 2×10^5^ cells were seeded on 12-well plates and grown to full confluence for 4 days. Then the cells were subjected to the first differentiation medium (DMEM supplemented 10% FBS, 0.5 mM 3-isobutyl-1-methylxanthine, 1 μM dexamethasone and 10 μg/ml insulin) starting on day 0 after confluence. After 2 days of induction, the medium was replaced with only insulin in DMEM with 10% FBS for an additional 2 days. Two days later, the cells were grown in regular medium for an additional 8 days and the medium was replaced every 2 days. Isobutyl-1-methylxanthine, dexamethasone and insulin were obtained from Sigma-Aldrich. 3T3-L1 cells were obtained from professor Yang of Nanjing Agriculture University. In this study, the medium was taken from 3T3-L1 adipocytes treated with spore varying from 10^6^ and 10^7^ CFU/ml for 24 h. Then cell culture supernatant from adipocyte was added to DCs for another 24 h.

### FTY720 treatments and tetramer staining

To inhibit circulation of memory T cells, FTY720 (Sigma) 1 mg/kg in PBS was administered intraperitoneally (i.p.), daily for 10 d. In addition, to assess the protective efficacy of the vaccines, the mice were immunized with the same vaccine. Intravascular staining was performed by injecting mice i.v. with FITC-conjugated anti-mouse CD45 antibody (5 μg) for 8-10 min before being euthanized. After immunization and FTY720 treatment, intestinal tissue was collected and collagenase digested, and cells were isolated for flow cytometric analysis as described previously [47]. The cells were stained with fluorochrome-conjugated antibodies or influenza HA-specific (HA_518–526_) H-2Kd tetramer-IYSTVASSL (MBL) reagent and analyzed using a flow cytometer (BD Biosciences). BMDCs (10^5^ cells) were incubated overnight in the presence of 10 μg/ml of influenza HA_518–526_ (IYSTVASSL) peptide (Genscript). IMALs were isolated from treated mice and A205804 was added to the plate for 5 d. The cells were stained with anti-CD3 for 20 min and HA_518–526_^+^ tetramer for an hour.

### DC/EC coculture system

DCs were generated from 4 to 6-week-old C57BL/6 mice using our previous method [48]. Briefly, bone marrow was extracted from the tibias and femurs of C57BL/6 mice with RPMI 1640. Then, the cells were suspended in complete medium (RPMI 1640 supplemented with 10% heat-inactivated FBS, 1% PenStrep), 10 ng/ml interleukin-4 (IL-4) and granulocyte-macrophage colony-stimulating factor (GM-CSF). After culture for approximately 60 h, the medium was lightly discarded to detach non-adherent granulocytes. Then clusters were harvested and subcultured overnight to remove adherent cells at 5 d. Non-adherent cells were collected at 6 d and used in subsequent studies. Caco-2 cells were seeded on the upper side of ThinCert membrane inserts (pore size, 3 μm) (Greiner Bio-One, Germany) in a 24-well plate overnight. The cells were maintained for 6 to 10 d until steady-state transepithelial electrical resistance of 300 Ω·cm2 was achieved [25]. In the coculture system, the filters were turned upside down, and then, DCs (5 ×10^5^ cells/ml) were cultured on the basolateral side of ECs for 4 h to let the cells attach to the filter. The filters were then turned right side up and placed into 24-well plates. The cells were incubated with different treatments with spore (10^7^ CFU/ml), or lipopolysaccharide (LPS) (1 μg/ml) for 24 h from the apical side. The filters and cells were fixed with 4% paraformaldehyde (PFA) for 15 min and processed for confocal microscopy. In addition, DCs were collected for phenotype assays and basolateral supernatants were collected for cytokine secretion assays.

### Ligated loop experiments

Mice were anesthetized with chloral hydrate (350 mg/kg body weight, intramuscular injection). The terminal ileal or jejunal ligated loop was injected with spore (10^8^ CFU/ml) or the same volume of PBS (0.01 M), and the intestines were removed after 0.5 h, optimal cutting temperature (OCT) (Tissue Freezing Medium, Sakura, Torrance, CA) and cut into 8 μm for immunofluorescence assays, as described below.

### Mouse cytokine array by a proteome profiler

Intestinal tissues from mice treated with PBS or spore for 7 d and 45 d were lysed with cell lysis buffer (R&D Systems) supplemented with 1% 0.2 mM phenylmethylsulfonyl fluoride (PMSF) at 4°C for 30 min. The protein concentration was detected with a protein bicinchoninic acid (BCA) kit (Thermo Fisher Scientific). Samples were analyzed with a mouse XL cytokine array kit (R&D Systems), according to the manufacturer’s instructions [49]. Immunospots were captured with an Odyssey Fc Imager (LI-COR), and data were analyzed with ImageJ software.

### Histology and immunohistochemistry

Immunohistochemistry detection was performed with the SABC kit (Boster Bioscience). Intrinsic peroxidase in samples was inactivated using 3% hydrogen peroxide after antigen retrieval was performed with buffer. Tissue sections were incubated with primary antibodies against ICAM-1 (1:200; Abcam) overnight at 4°C. Subsequently, the sections were incubated with biotinylated goat anti-mouse IgG as the secondary antibody. After staining with DAB, images were captured using a digital camera (Leica-DM4000B).

### Immunofluorescence (IF)

The fixed filters were permeabilized in 0.5% Triton X-100 in PBS for 5 min and blocked with 5% bovine serum albumin (BSA) in PBS for 2 h. Then the filters were stained with primary antibodies Armenian hamster anti-CD11c (N418) and rabbit anti-ICAM-1 (1A29, Abcam) overnight at 4°C, followed by incubation with secondary antibodies for 2 h at room temperature. For the *in vivo* model, cryosections were treated as described above. The filters were identified using confocal laser scanning microscopy (CLSM) (LSM 710; Zeiss, Oberkochen, Germany). Cross-sectional images were observed by z-axis views and analyzed using Zeiss ZEN 2012 and Adobe Photoshop CC (Adobe, San Jose, CA).

### Quantitative RT-PCR (qRT-PCR)

Total RNA from intestinal tissues was prepared using Trizol reagent (Takara, JPN) following the manufacturer’s guidelines and reverse-transcribed using a PrimeScript RT reagent Kit (Takara, JPN) according to the manufacturer’s instructions. QPCR was performed for triplicate samples using a SYBR Green qPCR Kit (Takara, JPN) by the Applied Biosystems™ QuantStudio™ 6 standard Real-Time PCR System (Thermo Fisher Scientific). The housekeeping genes β-actin was routinely used as internal controls. The primers used in this study were as follows: for β-actin, 5’-AAGTGTGACGTTGACATCCG-3’, rev 5’-GATCCACATCTGCTGGAAG-3’; for ICAM-1, 5’-TCACCAGGAATGTGTACCTGAC-3’, rev 5’-GGCTTGTCCCTTGAGTTTTATGG-3’.

### Western blot assay

The cells were lysed with RIPA buffer containing a 1% protease inhibitor cocktail on ice for 20 min. After removing debris by centrifugation at 4°C, supernatant protein was collected, and the total concentration was determined by a BCA protein assay kit. Protein was separated by electrophoresis on 10% sodium-dodecyl sulfate polyacrylamide gels (SDS-PAGE) and transferred to a polyvinylidene difluoride (PVDF) membrane. Mouse anti-ICAM-1 (1A29, Abcam), anti-Acrp30 (PA1-054, Thermo Fisher) and anti-β-actin (4D3, bioworld) were used to assess ICAM-1 and Acrp30 expression. Western blot images were visualized using an Image Reader Tanon-5200 imaging system.

### Statistical analysis

Results were shown as mean ± SEM. Student’s t-test was employed to determine that between two groups and One-way analysis of variance (ANOVA) with Dunnett’s test were performed with SPSS among multiple groups. The statistical analysis was performed using FlowJo v10, Microsoft Excel 2010 and Graph Pad Prism 7 Software. The asterisks indicate significant differences between H9N2 WIV plus spore and H9N2 WIV. P values < 0.05 were considered to be statistically significant (* *P* < 0.05, \*\**P* < 0.01).

## Acknowledgments

We thank for Penghao Zhang and Yuchen Li performed most of the immunization and in vitro immunogenicity experiments. We also thank for Jian Lin and Qinghua Yu conceived the study and revised the manuscript. Finally, we thank for Lulu Huang study conception and design, data analysis and interpretation, manuscript writing, final approval of the manuscript.

## Abbreviations

BMDCs: Bone marrow derived dendritic cells
AIV: avian influenza virus
Tcms: central memory T cells
Trms: Tissue-resident memory T cells
Tems: effector memory T cells
S1P: sphingosine-1-phosphate
TEDs: transepithelial dendrites
EC: epithelial cell
GM-CSF: granulocyte colony-stimulating factor
MHC-II: major histocompatibility complex class II
FACS: Fluorescence Activated Cell Sorter

## Key Resources Tables

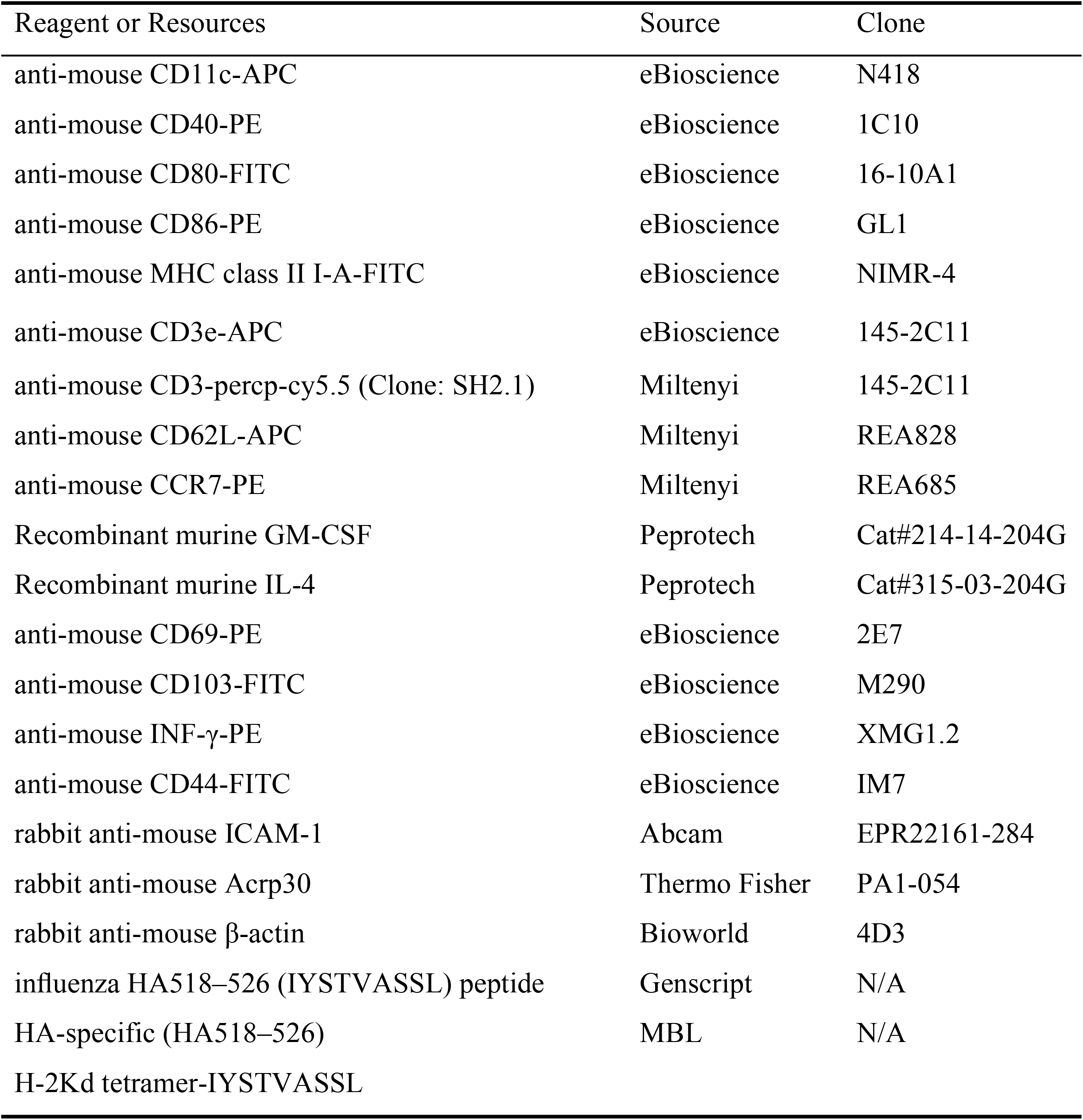

## Supporting information

**S1 Fig Spore plus H9N2 WIV generated abundant CD62L^+^CCR7^+^ cells during the early immunization period.**

The effect of spore on Tcms in the blood was detected by FACS. (A-D) The frequencies of Tcms (CD3^+^ CD62L^+^ CCR7^+^) were detected in the blood at 7 d (A, C) and 45 d (B, D) after priming immunization. A gating strategy of live cells and lymphocytes was applied, followed by gating for CD3^+^ cells and determination of the memory cell phenotype according to CCR7 and CD62L expression. The results are expressed as the mean ± SEM (n=6). \**P* < 0.05, \*\**P* < 0.01. One representative of three similar independent experiments is shown.

**S2 Fig Evaluation of tissue-resident T cells with a combination of FTY720 treatment and IV staining.**

(A, B) Vascular T cells were completely depleted by FTY720 treatment and stained by CD45-FITC I.V. staining. Schematic of vaccination, which was performed as described previously. Briefly, the immune-suppressive agent FTY720 (0.5 mg/kg/day) was administered intraperitoneally (i.p.) for 10 sequential days to deplete circulating lymphocytes at 6 weeks after the primary vaccination. On the following day, anti-CD45 mAb (FITC-labeled, 2.5 μg/mouse) was injected into the orbital vein to stain vascular leukocytes (IV staining) 10 min before euthanasia. Intestinal and peripheral blood leukocytes were collected to validate the efficacy of FTY720 administration and IV staining.

